# Evaluating restriction enzyme selection for genome reduction in conservation genomics

**DOI:** 10.1101/2022.11.26.518029

**Authors:** Ainhoa López, Carlos Carreras, Marta Pascual, Cinta Pegueroles

## Abstract

Conservation genomic studies in non-model organisms generally rely on genome reduction techniques based on restriction enzymes to identify population structure as well as candidate loci for local adaptation. These reduced libraries ensure a high density of SNP loci and high coverage for accurate genotyping. Despite the fraction of the genome that is sequenced is expected to be randomly located, the reduction of the genome might depend on the recognition site of the restriction enzyme used. Here, we evaluate the distribution and functional composition of loci obtained after Genotyping-by-sequencing (GBS) genome reduction with two widely used restriction enzymes (EcoT22I and ApeKI). To do so, we compared data from two endemic fish species (*Symphodus ocellatus* and *Symphodus tinca*, EcoT22I enzyme) and two ecosystem engineer sea urchins (*Paracentrotus lividus* and *Arbacia lixula*, ApeKI enzyme). In brief, we mapped the sequenced loci to the phylogenetically closest reference genome available (*Labrus bergylta* for fish and *Strongylocentrotus purpuratus* for sea urchins), classified them as exonic, intronic, and intergenic, and studied their functionality by using GO terms. We detected an enrichment towards exonic or intergenic regions depending on the restriction enzyme used, and we did not detect differences between total loci and candidate loci for adaptation. Despite most GO terms being shared between species, the analysis of their abundance showed differences between taxonomic groups, which may be attributed to differences of the targeted loci. Our results highlight the importance of restriction enzyme selection and the need for high-quality annotated genomes in conservation genomic studies.

## 1. INTRODUCTION

We are facing the sixth mass extinction on earth, with an accelerated global loss of biodiversity (IPBES, 2019). In the last decades, genetics has made it possible to delve into important processes of interest for conservation such as the level of inbreeding or gene flow between or within populations (Ouborg, Pertoldi, Loeschcke, Bijlsma, & Hedrick, 2010). However, there are still unresolved questions, and there is where conservation genomics plays a critical role. While conservation genetics is based on a reduced number of loci, conservation genomics is based on thousands of genome wide loci. Genomics improves our understanding of evolution and adaptation in the sea environment (Nielsen, Hemmer-Hansen, Larsen, & Bekkevold, 2009), even in marine non-model organisms. The usage of genome-wide loci allows the detection of adaptation patterns or population structure that were not detectable with fewer loci (Bradbury et al., 2015). High throughput genotyping techniques can be applied even in non-model organisms even without a reference genome (Andrews, Good, Miller, Luikart, & Hohenlohe, 2016). Among these techniques, genotyping-by-sequencing (GBS) is a simple system for building libraries for massive sequencing that allows the discovery of genome-wide SNP loci (Elshire et al., 2011). When using these techniques, it is assumed that the sequenced genome fraction is representative of the whole genome, but the reduction of the genome might depend on the recognition site of the restriction enzyme used. Consequently, the location of the candidate loci and their functional composition could be influenced by restriction enzymes, resulting in potential biases. To which extent the restriction enzyme selection influences genomic studies has not been evaluated yet.

Population genomic studies published in different taxa are of particular interest since they allow the evaluation of the effect of the genomic technique used (Carreras et al., 2020, 2021; Torrado, Carreras, Raventos, Macpherson, & Pascual, 2020). Carreras et al. studied the genetic structure of the two species of sea urchins cohabiting in the Mediterranean Sea: the edible sea urchin *Paracentrotus lividus* (2020) and the black sea urchin *Arbacia lixula* (2021). Sea urchins are important engineers of infralittoral benthic communities, playing a key ecological role controlling the structure of communities through grazing activity (Agnetta et al., 2015; Carreras et al., 2020; Palacín, Turon, Ballesteros, Giribet, & López, 1998; Wangensteen, Turon, & García-Cisneros, 2011). While *P. lividus* is mainly herbivorous, *A. lixula* has a tendency from omnivorous to carnivorous (Agnetta et al., 2013). Even though the two sea urchins have a role in the formation of barren patches (Bulleri, 2013; Bulleri, Benedetti-Cecchi, & Cinelli, 1999), some studies show that *A. lixula* has a role in maintaining them (Bonaviri, Vega Fernández, Fanelli, Badalamenti, & Gianguzza, 2011; Bulleri et al., 1999; Guidetti & Dulcić, 2007). Importantly, both species are facing the effects of global warming. The black sea urchin *A. lixula* is a thermophilic species (Pérez-Portela et al., 2019; Wangensteen, Turon, Pérez-Portela, & Palacín, 2012) contrary to the purple sea urchin *Paracentrotus lividus* that prefers cold waters. During the last years, populations of *P. lividus* have been declining, and some of them even collapsed mainly due to the high commercial interest (Yeruham, Rilov, Shpigel, & Abelson, 2015). In addition, the current increase in the sea water temperature is expected to favor *A. lixula*, due to its more thermophilic biology and its phenotypic plasticity (Pérez-Portela et al., 2019). In both species, Carreras et al. identified some degree of population structure and candidate loci for adaptation associated with salinity and different temperature variables. They mapped the small fraction of loci under selection, showing that numerous candidate loci were located in exonic regions, suggesting that candidate loci could be enriched at exonic regions (Carreras et al., 2020, 2021). The two endemic fishes from the Mediterranean sea *Symphodus ocellatus* and *Symphodus tinca* inhabit algal-covered rocky substrates and sea-grass beds like *Posidonia oceanica* (Macpherson, Gordoa, & García-Rubies, 2002). They are part of the Labridae family which represent a crucial connection of the trophic web in coastal environments (Shili, Souissi, & Bahri-Sfar, 2018). These two species are also considered supplementary fish cleaner, which help other fish (hosts) to be free of parasites (Zander & Sötje, 2002). While *S. ocellatus* is a microphagus predator (Macpherson et al., 2002), so it mainly feeds on Bryozoa, mollusks, and polychaetes (Quignard & Pras, 1986); *S. tinca* is a key species due to its abundance and generalist diet (Carreras et al., 2017), feeding on sea-urchins, ophioures and mollusks (Quignard & Pras, 1986). In addition to that, *S. ocellatus* can be used as a fish model for ecological impact studies due to its high density and distribution (Levi, Boutoute, & Mayzaud, 2005). Torrado et al. (2020) found different levels of population structure across the Western Mediterranean in the two species, with higher population differentiation in *S. ocellatus*. Nonetheless, they found several candidate loci in both species associated with temperature, productivity, and turbulence variables. However, most of the candidate loci identified by the authors were located in introns. In all four studies, loci were obtained by genotyping-by-sequencing (GBS) but using different restriction enzymes (ApekI for the sea urchins and EcoT22I for fish). The different composition of candidate loci may be attributed to differences between species, restriction enzymes, or biological adaptation. To answer this question, it is necessary to evaluate the composition of the candidate loci compared to all genotyped loci, which has not been addressed so far.

Here, we aim to test whether GBS data obtained using different restriction enzymes and species results in differential enrichment of genomic regions, functionalities or in both aspects, in total and candidate loci. To do so, we analyzed published data from four species (Carreras et al., 2020, 2021; Torrado et al., 2020), two endemic fish species obtained with the EcoT22I enzyme (*S. ocellatus* and *S. tinca*), and two ecosystem building sea urchins with the ApekI enzyme (*P. lividus* and *A. lixula*). We used the fasta files of all loci genotyped in the corresponding population genomic studies. First, we developed a pipeline that allows classifying loci as genic (distinguishing between exonic or intronic regions) and intergenic, using the closest available reference genome. Second, we evaluated the genomic composition of all annotated loci. We finally compared the genomic and functional composition of candidate loci and total loci, considering the different organisms and enzymes used.

## 2. MATERIALS AND METHODS

### 2.1 Species and data collection

We analyzed published population genomics data of two fish (Actinopterygii: *Symphodus ocellatus, Symphodus tinca*) and two sea urchins (Echinoidea: *Paracentrotus lividus* and *Arbacia lixula*) (Carreras et al., 2020, 2021; Torrado et al., 2020). Genomic loci for the four species were obtained by GBS using enzymes for genome reduction. The authors used EcoT22I for the two fish species, whose restriction site is (A | TGCA | T), where the bar identifies the cut sites generating sticky ends; and ApekI for the two sea urchins species, whose restriction site is (G | CWG | C) and W can be either A or T.

In fish (**Table S1**), we analyzed the 3,985 loci of *S. ocellatus* and 5,284 loci of *S. tinca* obtained using the STACKS v1.47 software in a previous study (Torrado et al., 2020). Loci were obtained from 162 individuals of *S. ocellatus* and 141 of *S. tinca* collected in 6 and 5 different locations respectively along the Mediterranean coast of the Iberian Peninsula. By using Redundancy analysis (RDA), Genome-wide association studies (GWAS), and Outlier analysis, the authors of this study identified a total of 292 and 168 loci to be under selection for *S. ocellatus* and *S. tinca* respectively (**Table S1**). In sea urchins (**Table S1**), we analyzed the 3,730 loci of *P. lividus* and the 5,241 loci of *A. lixula* obtained using the GIbPSs toolkit in Carreras et al. (2020, 2021). The loci were obtained from 241 individuals of *P. lividus* and 240 of *A. lixula* collected in 11 different locations from the occidental and oriental Mediterranean and the eastern Atlantic coast. By using RDA and Outlier analyses, the authors of these studies identified a total of 402 and 264 loci to be under selection for *P. lividus* and *A. lixula* respectively. We obtained the fasta files of the analyzed loci in the four species and used them in posterior analyses.

### 2.2 Classification and data analysis of total and candidate loci

To identify the genomic location of all the loci, we first mapped the sequences to the reference genome of the most closely related species using makeblastdb v2.10.1 followed by BLASTN searches (e-value ≤ 1e-4, outfmt = 6). In fish, we used the genome of *Labrus bergylta* (BallGen_V1, assembly accession: GCF_900080235.1) which diverged 28.2 MYA from *Symphodus ocellatus* and *Symphodus tinca* (http://www.timetree.org/ accessed in April 2022, **Figure 1a**). In sea urchins, we used the genome of *Strongylocentrotus purpuratus* as reference (Spur_5.0, assembly accession: GCF_000002235.5) which diverged 183 MYA from *A. lixula* and 53.9 MYA from *P. lividus* (http://www.timetree.org/ accessed in April 2022, **Figure 1a**). We then classified the significant blast hits into exonic, intronic, or intergenic regions using the in-house python scriptClassifyBlastOut.py (**Figure 1b**, script available in our GitHub repository (https://github.com/EvolutionaryGenetics-UB-CEAB/restrictionEnzimes.git). To calculate the percentage of exons, introns, and intergenic regions, we first converted the annotations of the genomes to BED12 format using gtfToGenePred, gff3ToGenePred, and genePredToBed scripts from USCS utils (http://hgdownload.cse.ucsc.edu/admin/exe/linux.x86_64/). Subsequently, we estimated the percentages using bedtools genomecov using -d and -split options (Quinlan & Hall, 2010).

**FIGURE 1:**
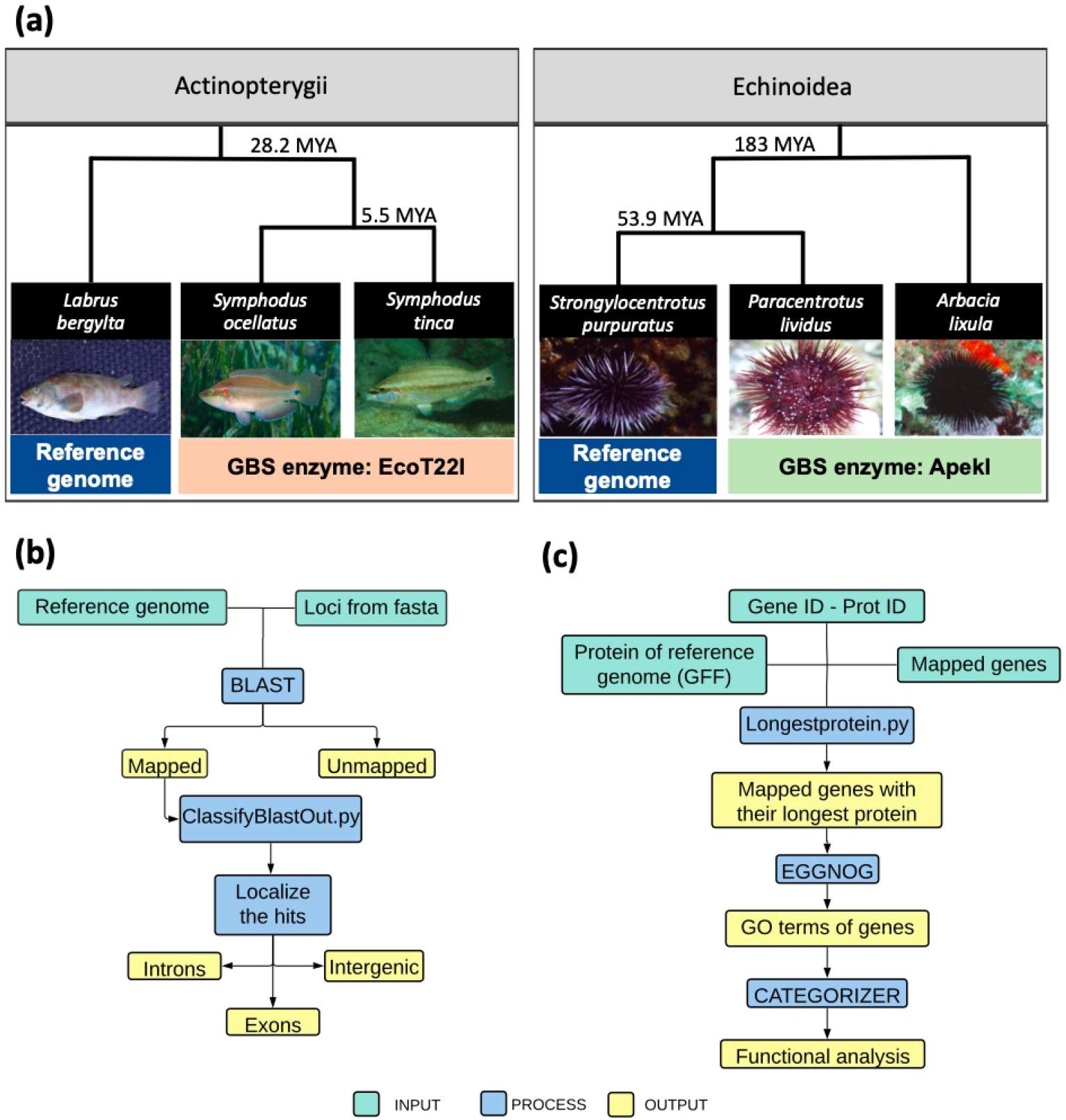
Workflow of the study design. (a) Representation of the data used in the study. We indicate the divergence time between the species from which we obtained GBS data (with the restriction enzyme used) and the reference genome used for each group of species. (b) Bioinformatic pipeline used to obtain the location of loci of each analyzed species to the reference genome. (c) Bioinformatic pipeline used to assign GO terms for the functional analysis. For each pipeline we detail the input data (cyan), the bioinformatic process involved (blue) and the output obtained in each analysis (yellow).

Count data were compiled in contingency tables. We checked for statistical differences in the loci classification within and between species by performing Fisher’s exact tests implemented in the package Stats 4.1.0 in R (R Core Team, 2021).

### 2.3 Functional analysis

For the functional analysis, we assigned GO terms to the loci with a positive hit of the four species based on the orthology relationship using eggNOG-mapper (Huerta-Cepas et al., 2019). To do so, we first made a list with the *L. bergylta* genes having significant blast hits with *S. ocellatus* and *S. tinca*, and a list with the *S. purpuratus* genes having significant blast hits with *P. lividus* and *A. lixula*. Using the GFF files from NCBI, we obtained the correspondence of the Gene ID and the Protein ID, and we obtained a fasta file with the longest amino-acid sequence for each identified gene. This file was used as input for the eggNOG-mapper using one-to-one orthology relationships within Metazoa. From the eggnog output file, we extracted the protein ID and the GO terms associated with them, and finally, we integrated the protein ID and GO terms with the genes name and locus ID of our species. **Figure 1c** shows a scheme of the pipeline used (https://github.com/EvolutionaryGenetics-UB-CEAB/restrictionEnzimes.git).

The analysis of the GO terms was done using the online server Categorizer (https://www.animalgenome.org/bioinfo/tools/countgo/, accessed 07/2022). First, we classified the GO terms according to the root category they belonged, to their biological process, to their molecular function, and to their cellular component. Secondly, we classified the GO terms assigned to the biological process category, by using the 442 categories from the GO slims list from QuickGO (https://www.ebi.ac.uk/QuickGO/). A Venn diagram showing the presence of GO terms in the GO slims categories for each of the 4 species was obtained using the ggvenn function from the ggplot2 package in R (Wickham, 2016). The visualization of the shared GO terms was performed using the Revigo software (Supek, Bošnjak, Škunca, & Šmuc, 2011). The counts data obtained from Categorizer was transformed into relative frequencies to be represented in a heatmap using the pheatmap package from R (Kolde, 2012).

## 3. RESULTS

### 3.1 Genomic characterization of total and candidate loci in fish and sea urchins

Overall, the frequency of loci mapping for each species to the respective reference genomes was low, with an average of less than 10% (**Figure 2a**). Within species, there were no significant differences in the mapping success for total and candidate loci (**Table 1**). We examined if there were significant differences between the two fish species (*S. ocellatus* vs. *S. tinca*) and between the two sea urchin species (*P. lividus* vs. *A. lixula*) for total and candidate loci mapping in the reference genomes (**Table S1**). The statistical tests showed no significant differences between *S. ocellatus* and *S. tinca* but significant differences between *P. lividus* and *A. lixula*, with a smaller frequency of mapped loci in the latter at both total and candidate loci (**Table 1**). Differences between taxonomic groups at total loci (fish vs. sea urchins) were significant when considering the four species, but non-significant differences arised when *A. lixula* was excluded from the statistical analysis (**Table 2**).

**FIGURE 2:**
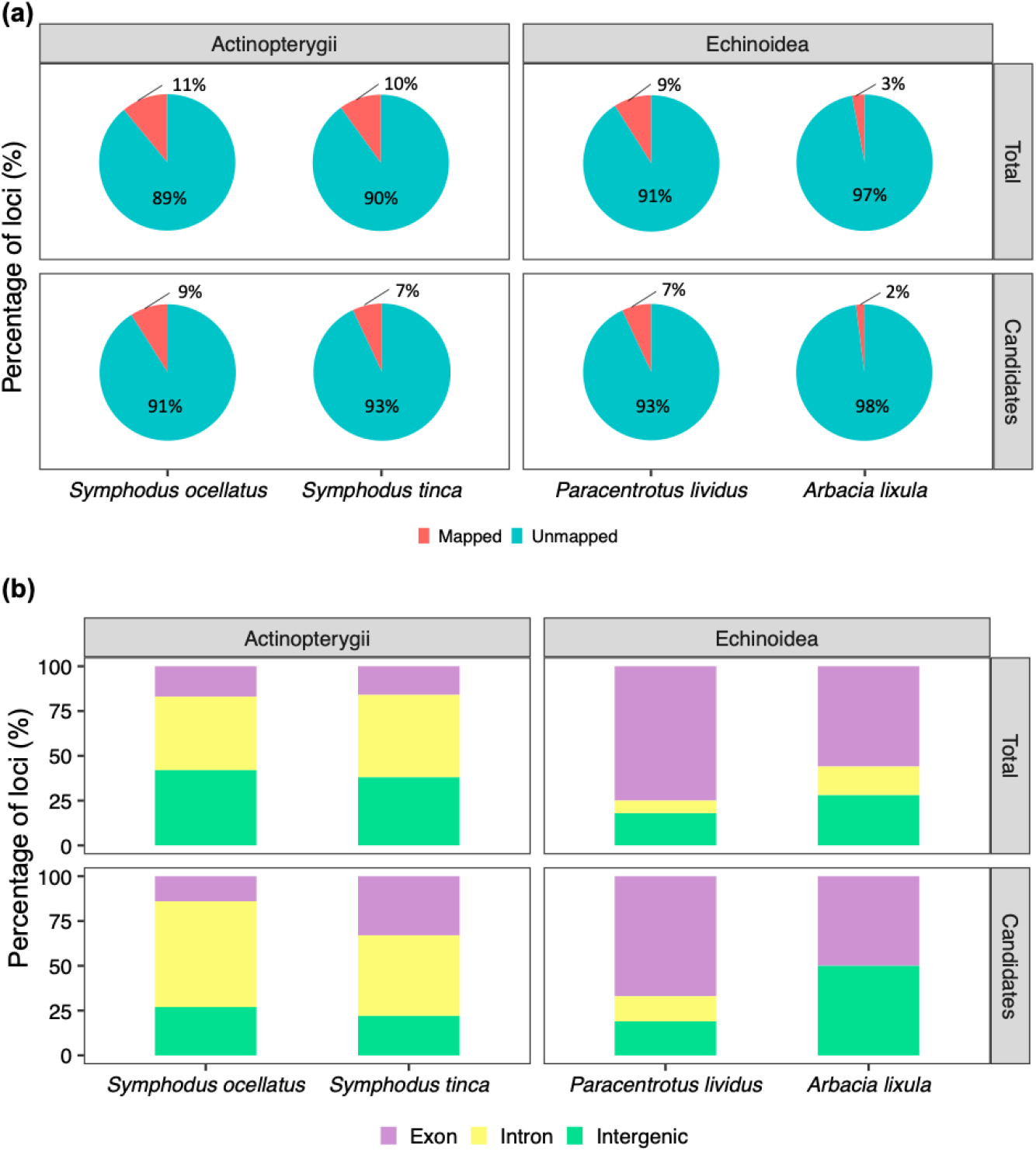
Mapping results of the GBS loci considered in this study. (a) Percentage of total (top) and candidate makers (bottom) that mapped to the closest reference genome for *S. ocellatus, S. tinca* (Actinopterygii), and *P. lividus* and *A. lixula* (Echinoidea). (b) Percentage of total and candidate mapped markers that were located in exonic, intronic, or intergenic regions for *S. ocellatus, S. tinca* (Actinopterygii), and *P. lividus* and *A. lixula* (Echinoidea).

**TABLE 1:** Fisher’s exact test p-value for the comparison between species and within species for the total and candidate loci classification. In bold are the significant values.

| Characteristics | Total vs. Candidate |  | *S. ocellatus* vs. *S. tinca* |  |
| --- | --- | --- | --- | --- |
|  | *S. ocellatus* | *S. tinca* | Total | Candidate |
| Mapped vs Unmapped | 0.428 | 0.229 | 0.144 | 0.477 |
| Unique vs Repeated | 1.000 | 1.000 | 0.666 | 1.000 |
| Exon vs Intron vs Intergenic | 0.288 | 0.31 | 0.349 | 0.492 |
| Characteristics | Total vs. Candidate |  | *P. lividus* vs. *A. lixula* |  |
|  | *P. lividus* | *A. lixula* | Total | Candidate |
| Mapped vs Unmapped | 0.312 | 0.476 | **0** | **0.004** |
| Unique vs Repeated | 1.000 | 0.682 | **0.008** | 0.161 |
| Exon vs Intron vs Intergenic | 0.335 | 1.000 | **0.001** | 0.585 |

**TABLE 2:** Chi-squared and p-value of total loci between sea urchins and fish including or excluding *A. lixula* in the comparison. In bold are the significant values.

| Characteristics | All species |  | Without *A. lixula* |  |
| --- | --- | --- | --- | --- |
|  | Chi-square | p-value | Chi-square | p-value |
| Mapped vs Unmapped | 221.49 | **<0.0001** | 4.70 | 0.094 |
| Unique vs Repeated | 88.96 | **<0.0001** | 21.50 | **<0.0001** |
| Exon vs Intron vs Intergenic | 328.46 | **<0.0001** | 312.27 | **<0.0001** |

The proportion of unique and repeated loci did not differ significantly between total and candidate loci for any of the species (**Table 1**). In fish, most loci (>82%) mapped to unique positions independently of being from the total or candidates loci set (Table S2). We did not detect significant differences between the two species of *Symphodus*, neither between total nor candidate loci (**Table 1**). In the case of *P. lividus* and *A. lixula*, we found significant differences for the total loci, with a higher frequency of unique loci in *P. lividus* **(Table S2)**. For candidate loci in sea urchins, differences were not significant, which may be due to the low number of candidate loci mapped, specifically two, in *A. lixula*. When we compared the frequency of unique loci between taxonomic groups, we obtained significant differences, with fish showing higher abundances independently of including or excluding *A. lixula* (**Table 2**).

We further classified the loci mapped to unique positions as exonic, intronic, or intergenic regions (**Figure 2b**). Overall, in the four species, we observed dominance of genic regions (exons and introns). However, the percentage of total loci mapping in genic regions was higher in *P. lividus* and *A. lixula* (82% and 72% respectively) than in *S. ocellatus* and *S. tinca* (59% and 62% respectively), despite the similar percentage of genic regions in their respective reference genomes (35%, **Table S3**). In fish, most loci in genic regions mapped to introns, contrary to sea urchins, where most loci mapped to exonic regions, despite the fact that the percentage of exons was very similar in the two reference genomes (6.5%, **Table S3**). There were no significant differences, between total and candidate loci within each species, in the frequency of loci mapping in exonic regions (**Table 1**). We did not detect significant differences between *Symphodus* species in the abundance of genes mapping in exonic regions in total or candidate loci, but we detected significant differences between sea urchins, specially when analyzing the total loci (**Table 1**). In addition, there were significant differences in exonic loci when comparing fish and sea urchins both considering and not considering *A. lixula* (**Table 2**).

### 3.2 Functional analyses

We performed a functional analyses in order to characterize the genes that were mapped uniquely to genes in the corresponding reference genomes, by assigning GO terms to the longest isoform using eggNOG mapper software. We assigned 9,196, 8,878, 4,146, and 4,310 GO terms to *S. ocellatus* (868 genes), *S. tinca* (162 genes), *P. lividus* (818 genes), and *A. lixula* (176 genes) respectively. All species had a similar percentage in the root classification of GO terms, where the most abundant was “biological process” including between 75-77% of the GO terms (Figure S2). We categorized the GO terms from the “biological process” category (6850, 6592, 3179, 3217 from *S. ocellatus, S.tinca, P. lividus*, and *A. lixula* respectively) using the 442 categories from GO slims. GO slims are a list of selected terms, including cytoplasm organization, metabolic process, DNA replication, localization, signaling, cell death, circadian rhythm, which help summarize GO terms into broad high-level categories. Using GO slims, we were able to classify 99% of the GO terms obtained from EggNogmapper. EggNogg mapper GO terms were assigned to 376 GO slims, representing 85.1% of the GO slims list terms. The number of counts obtained from the Categorizer was higher than the total number of input GO terms since one GO term can be classified in more than one category.

The majority of the GO slims (72.3%) were shared between the four species (**Figure 3a**). The GO slims shared by the four species were involved in a myriad of basic mechanisms, such as cellular component organization, metabolic process, development, localization, etc. (**Figure 3b, Table S4**). We also analyzed the relative abundance in each category by using a clustering analysis and the frequencies heatmap visualization (**Figure 3c**). Despite most GO slims having low frequency values, we detected two main clusters, separating fish and sea urchins.

**FIGURE 3:**
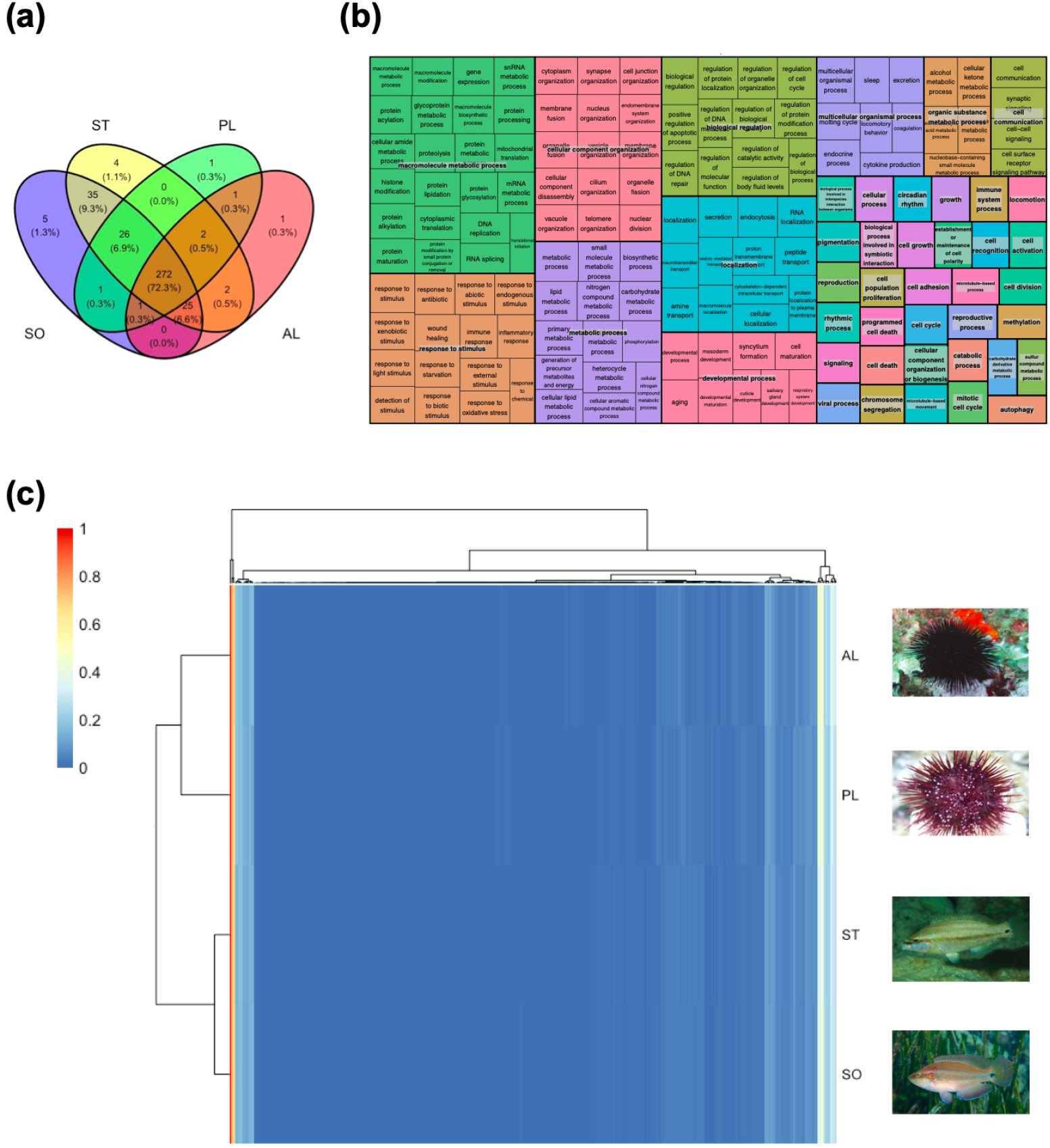
Results of the functional analysis. (a) Venn diagram showing the number of the classified GO slims terms in each species and shared among species. *SO: Symphodus ocellatus; ST: Symphodus tinca; PL: Paracentrotus lividus; AL: Arbacia lixula*. (b): Treemap of the 272 GO slims terms shared between all four species. The squares of similar functions are organized in the same color, with their representative GO term. (c) Heatmap of the 376 GO slims terms classified in Categorizer based on their normalized absolute frequencies. *SO: Symphodus ocellatus; ST: Symphodus tinca; PL: Paracentrotus lividus; AL: Arbacia lixula*.

## 4. DISCUSSION

Genomics is expected to revolutionize our understanding of the adaptive capabilities of endangered species and to improve their management strategies by ameliorating the delineation of their conservation units (CUs) (Funk, McKay, Hohenlohe, & Allendorf, 2012) Candidate loci for adaptation related to environmental cues are often identified in population genomic studies after using a genome reduction technique (Benestan et al., 2016; Sandoval-Castillo, Robinson, Hart, Strain, & Beheregaray, 2018; Torrado et al., 2022). The functional composition and gene category of candidate loci to be selected in several conditions and species have been studied in the past (Carreras et al., 2020; Pérez-Portela, Riesgo, Wangensteen, Palacín, & Turon, 2020; Schunter, Vollmer, Macpherson, & Pascual, 2014; Torrado et al., 2020). However, the distribution of all analyzed loci needs to be assessed to identify the processes leading to differences in genic distribution across studies and taxa. In the present work, we have shown that candidate loci obtained using the GBS technique are not enriched at certain genic categories but mirror the distribution of the total loci used in the population studies. However, their genic location may be greatly influenced by the methodology used, in terms of the nucleotide content of the recognition sequence of the restriction enzyme, as well as, the divergence time to the reference genome used for localizing the loci.

By mapping all loci to the closest reference genome, we observed that most loci were located in genic regions in the four species (69% of uniquely mapped reads on average in the four species, **Table S2**). Knowing that genes cover roughly one-third of the two reference genomes (**Table S3**), the enrichment towards genic regions suggests that the two restriction enzymes cut preferentially on genic regions. However, we detected significant differences when comparing the proportion of loci mapping to exons and introns between groups (sea urchins vs. fish). Sea urchins loci mostly mapped to exons, while fish loci mostly mapped to introns. Differences were significant despite including or not *A. lixula*, which was the species with the lowest fraction of mapped loci. It is worth noting that the genomic reduction technique used for obtaining the loci was the same for the four species (GBS) but the restriction enzyme used differed between groups: EcoT22I for fish, and ApekI for sea urchins. These two enzymes have two different restriction sites. The restriction site of EcoT22I is (A | TGCA | T), implying that the GC content is only 33% of the target. Conversely, the restriction site of ApekI is (G | CWG | C), implying that the GC content represents 80% of the target. Knowing that exons have a higher percentage of GC content compared to introns (Amit et al., 2012; Kalari et al., 2006), it is expected that the loci obtained after the ApekI enzyme (GC-rich) target a higher proportion of exonic regions, while the EcoTT22I enzyme (AT-rich) targets more non-exonic regions, such as introns and intergenic regions. These differences could not be attributed to differences in the genomes content, as the percentage of exons was very similar in the two reference genomes (6.5% of exons, **Table S3**). Thus, we can conclude that the differential enrichment towards intronic and exonic regions detected in fish and sea urchins respectively is due to the enzyme used in the genotyping.

Previous studies (DaCosta & Sorenson, 2014; Kirschner et al., 2016; Roszik, Fenyőfalvi, Halász, Karányi, & Székvölgyi, 2017) also reported a bias caused by the restriction enzymes, especially toward first exons. Thus, the assumption of sequencing random fractions of the genome is not accomplished, and it depends on the restriction enzyme selected. It is important to take this finding into consideration when designing a study for conservation purposes. For instance, conservation studies focusing on adaptation may benefit from GC-rich enzymes such as ApekI, while those focusing on neutral variability should select non-rich GC enzymes such as EcoT22I. However, it has been proposed that neutral and adaptive markers, which provide different types of information, should be integrated to make optimal management decisions to protect biodiversity (Funk et al., 2012).

A common objective of most studies based on genome reduction techniques besides looking at population structure is to identify loci having a significant role in adaptation (candidate loci). It can be expected that the genome composition of total and candidate loci may differ due to selective pressures on the candidate loci. For instance, the non-randomness of restriction sites could be due to the selective pressure for GC-rich restriction sites in coding regions (Bystrykh, 2013). However, when comparing the genomic composition of the total and candidate loci within species, we did not detect any significant difference for any of the four species analyzed, indicating that the candidate loci’s composition mirrors the total distribution. Further studies are needed to confirm this result since the number of mapped loci to the closest reference genome was low.

One of the striking results of our study is the low percentage of loci mapped to the closest available reference genome (less than 10%), likely a consequence of the divergence time between the reference genome species and the studied species. For instance, the percentage of loci mapped to the reference genome was higher in *P. lividus than A. lixula* (10% and 3% respectively), which is in agreement with their divergence time from *S. purpuratus* (58 MYA and 208 MYA respectively, **Figure 1**, **Figure S1**) Similarly in a previous study in fish, it was showed a significant negative correlation between the number of successfully microsatellite amplified and levels of polymorphism and the phylogenetic distance to the source species (Carreras-Carbonell, Macpherson, & Pascual, 2008). Furthermore, the number of reads mapped to its reference genome decreases according to the phylogenetic distance (Galla et al., 2018). Not only this, since genic regions are more conserved than the overall genome (Chaffey, 2003), the more phylogenetically distant the focus and reference species, the more likely to target genic regions, as we observed in the present study. Thus, the use of phylogenetically distant reference genomes plus the usage of GC-rich restriction enzymes will bias the results towards highly conserved genic regions, as we show in sea urchins. Despite the bias in genome category, the functional analysis showed that most of the functions assigned to the mapped loci were shared between the four species analyzed (**Figure 4, Table S4**). However, we detected several GO terms (7.2%) that were only assigned to fish. Since GO terms used in our analyses were involved in basic functions, their absence in sea urchins may be due to the different composition of genes sequenced in the two groups due to a differential evolutionary history. In agreement with this idea, when taking into account the frequency of the annotated GO terms, fish, and sea urchin loci cluster separately, suggesting functional differences between the loci of the two groups. Unfortunately, we could not perform a functional analysis of the candidate loci, due to the low percentage of loci mapped coupled with the lack of annotated GO terms in the reference genomes (annotations were transferred using orthology relationships). Altogether, conservation genomic studies based on genome reduction techniques will benefit from future high-quality and well-annotated reference genomes (Brandies, Peel, Hogg, & Belov (2019) and Formenti et al. (2019)). Luckily, their availability is increasing due to several initiatives such as the ERGA consortium or the Earth Biogenome Project (Formenti et al., 2022).

## 5. CONCLUDING REMARKS

This study demonstrates that the selection of the restriction enzyme is key when using genome reduction techniques in conservation genomics studies. We obtained compelling evidence that restriction enzymes produce important differences in the composition of mapped loci. Loci are biased towards exonic or intronic regions depending on the enzyme used. Although loci obtained are involved in a myriad of general functions, their functional composition seems to be affected by the loci targeted. The genome composition of candidate loci for adaptation mirrors one of the total loci in the four species analyzed. Importantly, we show that the number of loci mapped and characterized depends on the divergence time between the reference genome and the focal species, as well as, the reference genome quality. Our study highlights the need for well-annotated reference genomes for non-model species to dig deep into the functionality of the candidate loci identified in population genomic studies aiming at species conservation. In addition, it is critical to select the restriction enzyme according to the biological question that aims to be addressed.

## Supporting information

Table S4: number of loci associated to GO slims in each species

## AUTHOR CONTRIBUTIONS

All authors designed the research, analyzed the data, and contributed to writing the paper.

## ACKNOWLEDGEMENTS

This research was funded by MarGeCh (PID2020-118550RB, funded by MCIN/AEI/10.13039/501100011033) from the Spanish Government. The authors CC MP and CP are members of the research group SGR2017-1120.

## DATA ACCESSIBILITY STATEMENT

Genetic data were obtained from public repositories (*A. lixula*: PRJNA746276, *P. lividus*: PRJNA608661, *Symphodus ocellatus*: PRJNA646056 and *Symphodus tinca*: PRJNA646057). All the bioinformatic pipelines used in this research are available on GitHub (https://github.com/EvolutionaryGenetics-UB-CEAB/restrictionEnzimes.git).

## SUPPLEMENTARY INFORMATION

### Supplementary figures and tables

**FIGURE S1:**
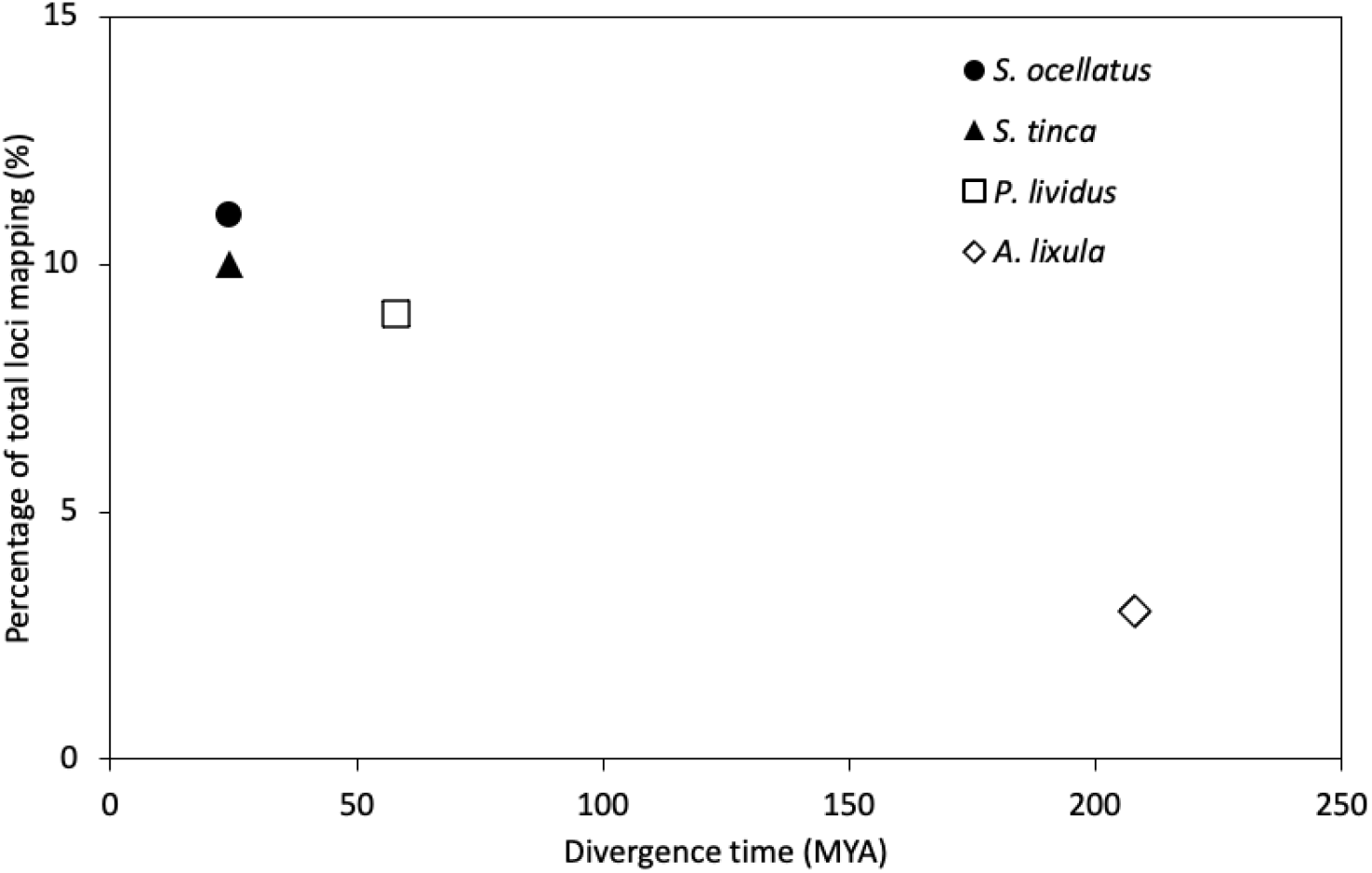
Percentage of total loci of the focal species mapping to the reference genome used and their divergence time in MYA. In fish (black symbols) the reference genome used was *Labrus bergylta;* in sea urchins (white symbols) the genome used as reference was *Strongylocentrotus purpuratus*. Each focal species has a different symbol.

**FIGURE S2:**
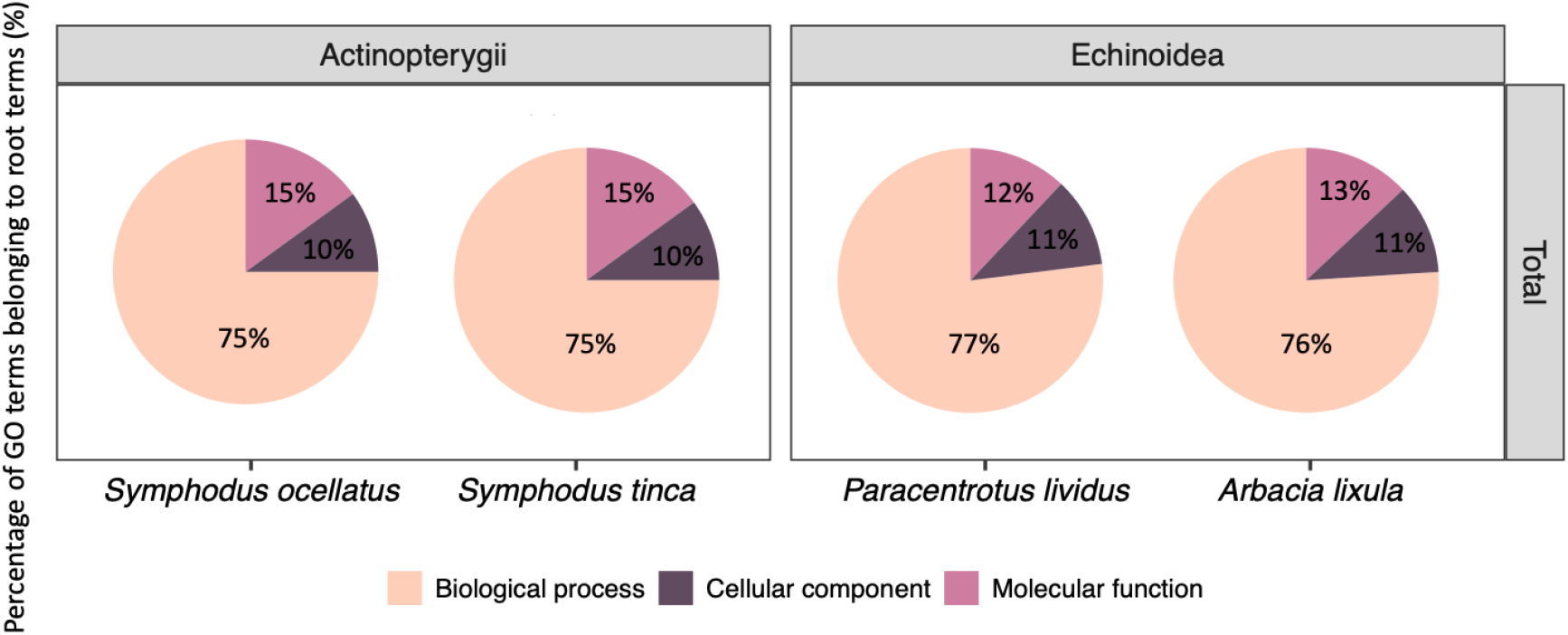
Classification of the GO terms for *S. ocellatus, S. tinca* (Actinopterygii), and *P. lividus* and *A. lixula* (Echinoidea) in root categories: Biological process, cellular component, and molecular function.

**TABLE S1:** For each species, grouped per taxon, we provide the number of individuals (N), and the total and candidate loci analyzed. Data from (Carreras et al., 2020, 2021; Torrado et al., 2020).

| Taxon | Species | N | Total loci | Candidate loci |
| --- | --- | --- | --- | --- |
| Actinopterygii | *Symphodus ocellatus* | 162 | 3985 | 292 |
|  | *Symphodus tinca* | 141 | 5284 | 168 |
| Echinoidea | *Paracentrotus lividus* | 241 | 3730 | 402 |
|  | *Arbacia lixula* | 240 | 5241 | 264 |

**TABLE S2:** Number of total and candidate loci identified in *S. ocellatus* (*So*), *S. tinca* (*St*), *P. lividus* (*Pl*) and *A. lixula* (*Al*) for the different analyzed characteristics in comparison to the corresponding reference genome.

| Characteristics | Total |  |  |  | Candidate |  |  |  |
| --- | --- | --- | --- | --- | --- | --- | --- | --- |
|  | SO | ST | PL | AL | SO | ST | PL | AL |
| Mapped | 423 | 512 | 342 | 174 | 26 | 11 | 30 | 6 |
| Unmapped | 3562 | 4772 | 3388 | 5067 | 266 | 157 | 372 | 258 |
| Unique | 352 | 420 | 242 | 88 | 22 | 9 | 21 | 2 |
| Repeated | 71 | 92 | 100 | 86 | 4 | 2 | 9 | 4 |
| Genic | 206 | 261 | 199 | 63 | 16 | 7 | 17 | 1 |
| Intergenic | 146 | 159 | 43 | 25 | 6 | 2 | 4 | 1 |
| Exon | 61 | 66 | 183 | 49 | 3 | 3 | 14 | 1 |
| Intron | 145 | 195 | 16 | 14 | 13 | 4 | 3 | 0 |

**TABLE S3:** Genome composition (in bp) of the two reference genomes.

|  | *Labrus bergylta* | *Strongylocentrotus purpuratus* |
| --- | --- | --- |
| **Genome** | GCF_900080235.1_BallGen_V1 | GCF_000002235.5_Spur_5.0 |
| **total** | 1,127,108,968 | 1,283,796,567 |
| **intergenic** | 732,365,203 (65.0%) | 838,232,936 (65.3%) |
| **genic** | 394,743,765 (35.0%) | 445,563,631 (34.7%) |
| **exons** | 73,115,318 (6.5%) | 83,622,857 (6.5%) |
| **introns** | 321,628,447 (28.5%) | 361,940,774 (28.2%) |

